# Decoding the microbiome and resistome of advanced chronic liver disease through long-read metagenomics

**DOI:** 10.64898/2025.12.09.693164

**Authors:** Hannah Trivett, Matthew J Dalby, Ned Peel, Darren Heavens, Raymond Kiu, Antia Acuna-Gonzalez, Merianne Mohamad, Gul Humayun, Richard M Leggett, Vishal C Patel, Lindsay J Hall

## Abstract

**Introduction:** Patients with advanced chronic liver disease (ACLD) and underlying cirrhosis frequently require repeated courses of antimicrobial therapy, with both the frequency and spectrum of antimicrobial exposure increasing alongside disease progression. In this population, impaired immune function and increased intestinal barrier dysfunction contribute to a heightened susceptibility of multidrug-resistant bacterial infections.

**Objective:** To comprehensively characterise the gastrointestinal microbiome and antimicrobial resistance gene (ARG) landscape across the clinical spectrum of ACLD, and to identify microbial and resistome signatures associated with disease severity.

**Design:** We employed long-read metagenomic sequencing (Oxford Nanopore Technologies) to profile the gastrointestinal microbiome and resistome across distinct ACLD stages: acute-on-chronic liver failure (ACLF), decompensated cirrhosis (DC), and stable cirrhosis, compared to healthy controls.

**Results:** ACLF patients had pronounced levels of *Enterococcus faecium*, with six samples out of 28 showing over 95% relative abundance, suggesting its potential as a bacterial biomarker for advanced cirrhosis. We reconstructed 28 high-quality MAGs of *E. faecium* from cirrhosis patients, 17 of which originated from ACLF cases. This dominance of *E. faecium* correlated with substantially reduced microbial diversity and a marked depletion of key commensal taxa, including *Blautia, Akkermansia, Faecalibacterium,* and *Bifidobacterium*. Resistome analysis revealed significant enrichment of clinically relevant ARGs in DC and ACLF, including those conferring resistance to aminoglycosides, beta-lactams, and glycopeptides, correlating with prior antimicrobial exposure.

**Conclusion:** Long-read metagenomics enables high-resolution characterisation of microbial and resistome dynamics across ACLD severities. By capturing taxonomic shifts, functional potential, and ARG enrichment, this approach provides valuable insights into microbiome trajectories linked with disease severity, informing mechanistic research and potential clinical interventions.

**Impact statement:** Chronic liver disease is a major global health burden, responsible for approximately 2 million deaths each year. Advanced chronic liver disease profoundly disrupts the gut microbiome, often exacerbated by repeated antibiotic exposure, promoting the persistence of antimicrobial-resistant organisms. Leveraging Oxford Nanopore Technologies long-read metagenomic sequencing, this study delivers high-resolution insights into the gut microbiome and resistome across progressive stages of cirrhosis. We reveal *Enterococcus faecium* dominance as a defining feature of cirrhosis, accompanied by severe loss of commensal diversity and enrichment of clinically significant resistance genes. These findings underscore the clinical utility of culture-independent metagenomic profiling for detecting pathogenic taxa and resistance determinants within the gut ecosystem of high-risk patients. Our work highlights the translational potential of long-read metagenomics as an accessible tool for pathogen surveillance, antimicrobial stewardship, and precision infection management in advanced liver disease.

## Background

Advanced chronic liver disease (ACLD) represents a major global health burden, encompassing a spectrum from compensated cirrhosis to decompensated cirrhosis (DC) and acute-on-chronic liver failure (ACLF). These stages are associated with high morbidity and mortality, with ACLF having a high short-term mortality of 90 days in over 50% of patients (Mahmud *et al*., 2019; Trebicka, Fernandez, *et al*., 2021).

Complex interactions between host factors, microbiome perturbations, and systemic inflammation drive disease progression in ACLD and increase susceptibility to bacterial infections (Miranda-Zazueta *et al*., 2020). In cirrhosis, disruption of the intestinal barrier facilitates bacterial translocation from the gut into ascitic fluid and systemic circulation, leading to complications such as sepsis and spontaneous bacterial peritonitis (Woodhouse, Singanayagam and Patel, 2020).

Emerging evidence indicates that the gut microbiome plays a central role in the pathophysiology of ACLD, influencing immune regulation, intestinal barrier integrity, and susceptibility to infections (Trebicka, Macnaughtan, *et al*., 2021; Jin *et al*., 2025). Progressive disease stages are characterised by microbiome perturbations, including the depletion of beneficial commensals such as *Faecalibacterium, Ruminococcus,* and *Bacteroides,* alongside reduced levels of key metabolites, including short-chain fatty acids, that maintain epithelial integrity and gut permeability. These shifts coincide with the expansion of potentially pathogenic taxa, i.e. pathobionts, that can dominate the gastrointestinal tract, including *Enterococcus faecium, Streptococcus thermophilus*, and *Ruminococcus lactaris* (Solé *et al*., 2021). In patients with ACLD, increasing overlap of the oral and gut microbiomes with progressive disease severity spanning SC, DC and ACLF is accompanied by a marked expansion of virulence-factor and antimicrobial resistance gene (ARG) carriage (oral >1,200 ARGs; gut >670 ARGs), highlighting resistome evolution in cirrhosis-associated perturbations (Lee *et al*., 2025). However, most existing studies have relied on short-read or targeted sequencing approaches, limiting resolution of microbial genomes and their functional attributes, including ARGs.

Antimicrobial resistance is a critical concern in ACLD, where frequent antibiotic use for prophylaxis and infection management may select for multidrug-resistant organisms (Patel and Williams, 2020). It is estimated that approximately one-third of bacterial infections in patients with cirrhosis are caused by multidrug-resistant bacteria, resulting in a fourfold increase in mortality risk compared to infections caused by non-resistant bacteria (Terra *et al*., 2023). In clinical settings, culture-based pathogen detection remains the standard practice; however, these methods are limited by slow turnaround times, poor detection of fastidious species, and low throughput, often resulting in the missed detection of polymicrobial infections (Chen *et al*., 2022; Hassall *et al*., 2024). The increasing adoption of multiplex PCR has improved diagnostic speed; however, as these assays rely on predefined primer targets, they are restricted to known organisms. Consequently, both culture-based and molecular methods may fail to identify clinically relevant or novel pathogens and overlook broader genomic context, such as gene function, AMR, and secondary microbial contributions to infections (Batool and Galloway-Peña, 2023).

Metagenomic sequencing, particularly long-read technologies, offers unique advantages for resolving complex microbial communities and their potential metabolic pathways, virulence factors, and resistome. Crucially, it enables the recovery of high-quality metagenome-assembled genomes (MAGs) and accurate ARG profiling, which conventional diagnostic methods can miss. Applying this approach to ACLD could uncover microbial signatures and resistance determinants associated with disease severity, providing insights into host-microbe and microbe-microbe interactions that underpin clinical outcomes (Shi *et al*., 2024). Indeed, with the appropriate infrastructure, metagenomics can yield actionable results with a rapid turnaround from sample collection to pathogen identification in less than 24 hours (Gu *et al*., 2021; Alcolea-Medina *et al*., 2024; Sung *et al*., 2025).

In this study, Oxford Nanopore Technologies (ONT) long-read metagenomics was employed to characterise the gut microbiota and resistome across the clinical spectrum of ACLD - stable cirrhosis, DC, and ACLF, comparing them to healthy controls. We aimed to identify taxonomic profiles and functional changes associated with ACLF progression, with a focus on MAGs and ARG enrichment profiles. Our findings show that liver disease progression is marked by pronounced gut microbiota disruptions and enrichment of ARGs, highlighting distinct microbial and resistome signatures that could inform clinical monitoring and therapeutic strategies.

## Methods

### Study participants & biological sampling

Patient participants, or their family nominee as consultees in the case of lack of capacity, provided written informed consent within 48 hours of presentation. Patients were managed according to standard evidence-based protocols and guidelines (Angeli *et al*., 2018). Main exclusion criteria included pregnancy, hepatic or non-hepatic malignancy, pre-existing immunosuppressive states, replicating hepatitis B or C or HIV infection, and known inflammatory bowel disease.

Demographic, clinical, and biochemical metadata were collected at the time of biological sampling. Standard clinical composite scores used for risk stratification and prognostication included the Child-Pugh score (Pugh *et al*., 1973), model for end-stage liver disease (MELD) (Kim *et al*., 2008), United Kingdom model for end-stage liver disease (UKELD) (Asrani and Kim, 2010), Chronic Liver Failure Consortium-acute decompensation (CLIF-C AD).

Healthy controls (HCs) aged >18 years were recruited to establish reference values for the various assays performed. Exclusion criteria for healthy controls were body mass index <18 or >27; pregnancy or active breastfeeding, a personal history of thrombotic or liver disease; chronic medical conditions requiring regular primary or secondary care review such as inflammatory bowel disease, and/or prescribed pharmacotherapies.

### Faecal sample acquisition

Faecal samples were obtained within 48 hours of admission to the hospital and collected into non-treated sterile universal tubes (Alpha Laboratories™), without any additives. Faecal samples were kept at 4°C without any preservative and, within 2 hours, were homogenised, pre-weighed into 200mg aliquots in FastPrep tubes (MP Biomedicals™), for storage at −80°C for subsequent DNA extraction.

### DNA extraction and long-read metagenomic sequencing

The FastDNA Spin Kit for Soil (MP Biomedicals™) was used to extract DNA from faecal samples, following the manufacturer’s instructions, with an extended one-minute bead-beating using the FastPrep-24 (MP Biomedicals™), with a speed setting of 6.0. DNA concentrations of each sample were determined using the Qubit Broad Range assay (Life Technologies), and molecule length was analysed on an Agilent TapeStation using Genomic Tape and reagents (Agilent Technologies).

Samples were pooled, with up to 400 ng of DNA input per sample, using the SQK-LSK109.24 ONT kit, which accommodates up to 18 samples in a single pool. All sequencing was performed on MinION R9.4.1 flowcells on an ONT GridION sequencer.

### Metagenomic read profiling using MARTi

ONT sequencing data in FAST5 format was converted to POD5 format using POD5 v0.3.6 (https://github.com/nanoporetech/pod5-file-format). Basecalling and demultiplexing of the raw signal data were performed using Dorado v0.8.3 with the dna_r9.4.1_e8_sup@v3.6 model (https://github.com/nanoporetech/dorado). Taxonomic classification and AMR gene detection were performed with MARTi v0.9.28 (Peel *et al*., 2025) on the first 100,000 basecalled reads per sample that passed the prefiltering criteria, minimum read length of 150 bp and minimum quality score of eight. For taxonomic classification, we used MARTi’s BLAST-based Lowest Common Ancestor pipeline against NCBI’s core nucleotide (core_nt) database (October 2024 release). Where specified, an LCA threshold of 0.1% was used – this means that any taxa representing fewer than 0.1% of the classified reads will be moved up a taxonomic level (e.g. from species to genus) until the threshold is reached. AMR gene detection was performed using the CARD database (Alcock *et al*., 2023) v4.0.0 (nucleotide_fasta_protein_homolog_model.fasta). Resulting taxonomic and AMR classifications were explored and visualised using the MARTi graphical user interface.

### Sequencing read quality control and metagenomic assembly

Long read metagenome raw reads (FAST5) were converted into POD5 files using POD5 v0.3.15 (https://github.com/nanoporetech/pod5-file-format). The converted files for each sequencing run were basecalled and demultiplexed using Dorado v0.8.3 (https://github.com/nanoporetech/dorado) with the selected model dna_r9.4.1_e81_sup@v3.6. Next, raw FASTQ reads produced by Dorado were trimmed and quality-filtered (q15) using fastplong (v0.2.2) (Chen, 2023). Subsequently, host-associated contamination (human reads) was removed using Hostile v2.0.0 (Constantinides, Hunt and Crook, 2023) under default parameters.

Cleaned metagenome reads were assembled using metaFlye (v2.9.5) (Kolmogorov *et al*., 2020). The assembled contigs then underwent three rounds of polishing using Medaka v2.0.0 (https://github.com/nanoporetech/medaka) with the model r941_e81_sup_g514.

### Metagenome-Assembled Genome binning

The MetaWRAP pipeline (v1.3) (Uritskiy, DiRuggiero and Taylor, 2018) generated metagenome bins. Bins were generated using three binning software tools within the binning module: CONCOCT (v1.1.0) (Alneberg *et al*., 2014), MetaBAT (v2.12.1) (Kang *et al*., 2015, 2019), and MaxBin (v2.2.6) (Wu *et al*., 2014). The bins were refined to generate a final set of bins using the bin refinement module within MetaWRAP, based on quality assessment using CheckM (v1.1.3) (Parks *et al*., 2015)with parameters of>70% completeness and <10% contamination. Metagenome bins were characterised using GTDB-Tk (v2.4.0) via the classify_wf module (Chaumeil *et al*., 2022). Following binning with MetaWRAP, three samples were excluded due to the absence of high-quality bins, as defined by CheckM quality metrics (completeness ≥ 70% and contamination ≤ 10%).

In total, 389 bins were produced across the metagenomes, including 27 bins identified as *E. faecium.* Of the 389 bins, two bins were characterised as archaea, specifically *Methanobrevibacter smithii*. All MAGs were dereplicated using dRep (v3.4.2) (Olm *et al*., 2017) with an ANI 99.9% as the strain-level inference cut-off, producing 382 dereplicated MAGs.

A neighbour-joining tree was produced for the dereplicated genomes using Mashtree (v1.4.6) (Katz *et al*., 2019) with 1000 bootstrap replicates and --min_depth 0. The tree was then mid-point rooted and visualised in iTOL (v7.2) (Letunic and Bork, 2024a).

### Taxonomic profiling

Following the quality filtering steps, taxonomic profiling of the filtered metagenomic reads (in FASTQ format) was performed using Sourmash (version 4.8.12) (Irber *et al*., 2024). Analyses were conducted with a k-mer size of 51, utilising the GTDB-rs214 database. The workflow incorporated the sketch, gather, and tax annotate commands for taxonomic classification, enabling the estimation of taxa across the metagenomes. Using R (V4.4.2) (Vienna, Austria: R Foundation for Statistical Computing, 2025), within RStudio (version 2025.05.1) (Posit team, 2025), the taxonomy-annotated output was converted into a Phyloseq object using Sourmashconsumr (Chou and Reiter, 2024) (v0.1.0). Phyloseq (McMurdie and Holmes, 2013) (v1.50.0) was utilised to calculate diversity metrics, including the relative abundance of taxa and alpha diversity. This was visualised using ggplot2 (Villanueva and Chen, 2019).

### AMR profiling of the metagenome

Antimicrobial resistance gene profiles were generated for the assembled metagenomes using AMR Finder-Plus (v4.0.19) with the database version 2024-12-18.1 to identify antimicrobial resistance genes, employing the parameters --ident_min 0.95 and --coverage_min 0.9 (Feldgarden *et al*., 2021). The resistome analysis of the metagenome was performed on the polished metagenome assemblies, whilst individual bacterial genome profiling was performed on the MAGs obtained via MetaWRAP (Uritskiy, DiRuggiero and Taylor, 2018).

Statistical analyses were conducted in *R* (version 4.4.2) (Vienna, Austria: R Foundation for Statistical Computing, 2025) using RStudio (version 2025.05.1) (Posit team, 2025). Faecal antimicrobial resistance (AMR) gene data were analysed as presence/absence matrices. Community-level differences were explored using non-metric multidimensional scaling (NMDS) based on Jaccard dissimilarities, implemented in vegan (v2.6-10) (Oksanen *et al*., 2001). Assumptions of homogeneity of group dispersions were evaluated with the betadisper function and found to be significantly different. To account for this, group differences were tested using an analysis of similarity (ANOSIM) with 999 permutations and Jaccard distances. Pairwise ANOSIM was used to assess differences between individual groups, with false discovery rate (FDR) correction for multiple testing.

Differences in the presence/absence of individual AMR genes across the four groups were assessed with Fisher’s exact test, or Chi-squared tests where all cell counts exceeded five. Genes with fewer than 10 total occurrences across all samples were excluded.

Differential relative abundance of bacterial species between groups was tested using the Kruskal–Wallis rank-sum test, appropriate due to the absence of some abundant species from certain participant groups. Relative abundances were compared across groups for each species, with significant results (P < 0.05) followed by post-hoc pairwise Dunn’s tests with FDR correction. Kruskal–Wallis P values across all taxa were also corrected using FDR.

For all analyses, statistical significance was defined as P < 0.05 after adjustment for multiple testing where applicable.

### Plasmid contig identification and AMR profiling

Metagenomic contigs were characterised as plasmid, virus or chromosome using GeNomad (v1.11.0) (Camargo *et al*., 2024) using the end-to-end classification module. Plasmid assigned contigs were profiled using AMR Finder-Plus (v4.0.19) (Feldgarden *et al*., 2021) as described above, to predict AMR genes present on plasmid contigs. In addition, plasmid identification was predicted using ABRicate (v1.0.1) (https://github.com/tseemann/abricate) via PLSDB (v2024_05_31_v2) (Galata *et al*., 2019) with options --minid=90 and --mincov=90.

### Enterococcus faecium profiling

Those classified as *E. faecium* were oriented using Circulator (v1.5.5) to orient genomes to begin at *dnaA* (Hunt *et al*., 2015). The average nucleotide identity was calculated between the *E. faecium* MAGs using FastANI (v1.34) using default parameters (Jain *et al*., 2018). Additionally, antimicrobial resistance gene profiles were generated using AMR Finder-Plus (v4.0.19) as described above.

For comparative purposes, 22 *E. faecium* genome sequences were retrieved from NCBI from Lebreton et al (Lebreton *et al*., 2013), as representatives of clade A1 (n=11) and clade A2 (n=11). MLST was predicted using PubMLST, under default parameters against the *E. faecium* typing scheme, providing allelic profiles, sequence type and estimations of clonal complexes for the *E. faecium* MAGS and reference genomes (Jolley, Bray and Maiden, 2018).

### Phylogenetic core gene SNP tree generation

The reference genomes along with the MAGs were then annotated using Bakta (v1.11.0) under default parameters using the full version 6 database, which consists of 10 publicly available reference databases (https://zenodo.org/records/14916843) (Schwengers *et al*., 2021). A core gene alignment was generated using Panaroo (v1.5.2) (Tonkin-Hill *et al*., 2020), which merged paralogs, while SNP variants were extracted with snp-sites (v2.5.1) (Page *et al*., 2016). A phylogenetic tree was generated using IQ-TREE (Minh *et al*., 2020). ModelFinder (Kalyaanamoorthy *et al*., 2017) within IQTREE was employed to find the best-fitting model for the species, selecting the GTR+F+I+R6 model to construct the phylogenetic tree using 1000 ultrafast bootstrap replicates (Hoang *et al*., 2018). ClonalFrameML (v1.13) was employed to account for recombination when inferring the phylogenetic relationships among the genomes (Didelot and Wilson, 2015). iTOL (v7.2) was used for tree visualisation and annotation (Letunic and Bork, 2024b).

## Results

### Baseline demographics and clinical characteristics

This cross-sectional study included 76 participants categorised into four cohorts: healthy controls (HC), stable cirrhosis (SC), decompensated cirrhosis (DC), and acute-on-chronic liver failure (ACLF). Of these, 32 participants identified as female and 44 as male. The HC group comprised 12 hospital staff members and served as a reference for comparative analyses and establishing baseline values. Among the 64 participants diagnosed with ACLD, the distribution by stage was as follows: SC (n = 20), DC (n = 16), and ACLF (n = 28). ACLD aetiology spanned 17 distinct diagnostic categories, with the most common being alcohol-related liver disease (ALD) in abstinent individuals (n=26), ALD in active drinkers (n=8), and metabolic dysfunction–associated steatotic liver disease (MASLD), previously known as non-alcoholic fatty liver disease (n=7). The mean age for the HC cohort was 36 years, compared to 53 years for patients with cirrhosis overall (SC = 56 years, DC = 52 years, ACLF = 49 years). Antimicrobial exposure was frequent across the ACLD cohort, with 62.5% receiving treatment as part of standard clinical infection management regimens. ACLF patients, as expected, had the highest exposure, with 27 of 28 patients (96.4%) receiving antimicrobials, often as combination therapy. Fourteen ACLF patients were treated with two or more agents, including antibacterial and antifungal combinations (Supplementary Table 1).

### Progression of ACLD is associated with reduced taxonomic diversity

Alpha diversity showed a progressive decline across ACLD severity groups. The Chao1 index, which estimates species richness, was highest in the HC cohort and decreased sequentially in the SC, DC, and ACLF groups (Figure 1A). Similarly, the Shannon index, which accounts for both richness and evenness, revealed a stepwise reduction in microbial diversity, with the lowest values observed in ACLF patients (Figure 1B). The Inverse Simpson diversity index indicated a significant difference in microbial diversity between HC and each disease state. A Kruskal-Wallis test confirmed significant differences in microbial diversity across groups (χ² = 48.13, df = 3, p < 0.001). Pairwise Wilcoxon tests with the Benjamini-Hochberg correction showed all group comparisons were significant (p < 0.001), suggesting a progressive decline in diversity from HC to ACLF (Figure 1C). Non-metric multidimensional scaling (NMDS) showed that SC clustered closely with HC, while DC had an intermediate position between SC and ACLF, exhibiting partial overlap with ACLF. PERMANOVA analysis based on Bray-Curtis dissimilarity revealed significant differences in microbial community composition among groups (R² = 0.08, p = 0.001) (Figure 1D).

**Figure 1.**
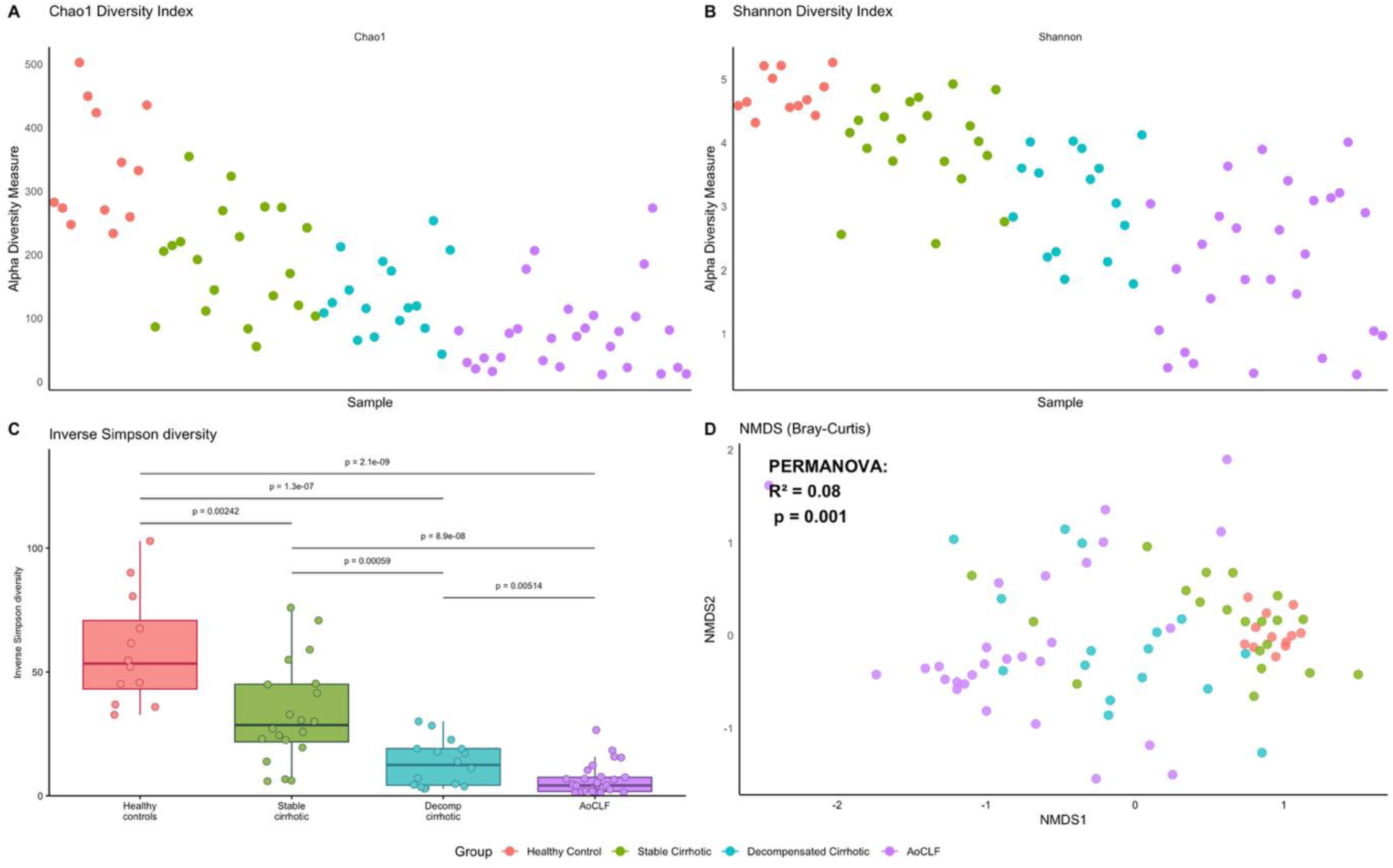
Comparison of microbial diversity profiles of faecal samples from healthy controls and cirrhosis patients. **A)** Chao1 diversity index representing species richness. **B)** Shannon diversity index accounting for species richness and evenness. **C)** Inverse Simpson’s Index measuring microbial diversity across cohorts, post hoc pairwise Wilcoxon tests (Benjamini-Hochberg adjusted) revealed significant differences between the HC cohort and each disease state. **D)** NMDS plot based on Bray-Curtis dissimilarity with PERMANOVA significance (R² = 0.80, p = 0.001) overlaid.

The gut microbiome differed significantly between HC and ACLD cohorts when comparing bacterial diversity using taxonomic profiling (Figure 2A). Stacked bar plots revealed a progressive decline in the relative abundance of beneficial bacterial genera, including *Anaerostipes*, *Agathobacter*, *Bifidobacterium*, *Blautia*, *Faecalibacterium*, *Fusicatenibacter*, and *Ruminococcus.* In parallel, an increase in pathogenic and opportunistic taxa was observed, including *Enterococcus*, *Escherichia*, and *Klebsiella*. These changes were more pronounced in ACLF, where *Enterococcus* dominated microbial communities. Detailed species-level analysis (Figure 2B) showed that HC samples species were significantly more diverse than those of cirrhosis patients, characterised by several keystone taxa such as *Faecalibacterium prausnitzii*, *Anaerostipes hadrus*, *Agathobacter rectalis*, *Bifidobacterium longum*, *Blautia_A wexlerae and Ruminococcus_E bromii_B*. Post hoc pairwise comparisons using Dunn’s test revealed significant differences between the HC and ACLD cohorts for *A. rectalis, B. wexlerae*, and *F. saccharivorans*, with their abundances progressively declining in samples as cirrhosis severity increased (Figures 2C, 2D, and 2E). Conversely, pathogenic bacteria dominated the gastrointestinal microbiome in the later stages of liver disease, replacing beneficial taxa, most notably *Enterococcus faecium*, *Escherichia coli*, and *Klebsiella pneumoniae*. The *Enterococcus* genus was absent in HC but showed a stepwise increase in both relative abundance and prevalence with advancing disease severity, particularly *E. faecium*. In the ACLF cohort, *E. faecium* was present in the majority of samples, with 10 individuals exhibiting a relative abundance of more than 50%. This marked overrepresentation underscores the shift towards a pathogen-enriched community and the potential clinical relevance of *E. faecium* overgrowth in ACLD progression. Differential abundance testing using the Kruskal-Wallis test revealed significant differences in *E. faecium* abundance among study groups (FDR-adjusted p-value < 0.001). Post hoc Dunn’s tests showed *Enterococcus* was significantly enriched in ACLF compared to HC, SC, and DC; however, no significant difference was found between HC and SC (FDR-adjusted p-value = 0.267) (Figure 2F).

**Figure 2.**
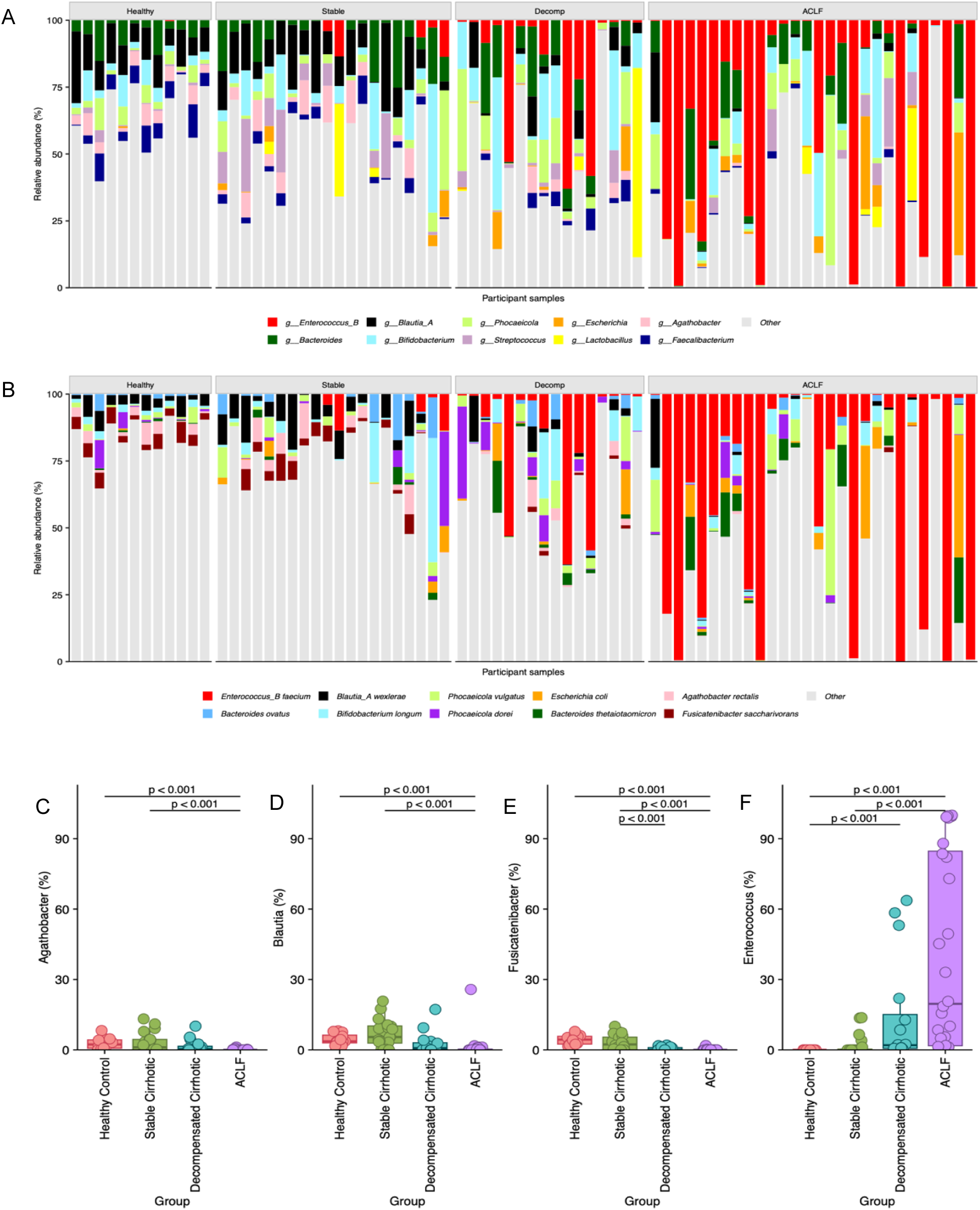
Microbial composition in cirrhosis patients compared to healthy controls. **A)** Stacked bar plot of the relative abundance of the top 10 bacterial genera across cohorts, ordered by disease severity. **B)** Stacked bar plot of the top 10 bacterial species present across cohorts, ordered by disease severity. **C-F) Box plots showing relative abundance of species significantly enriched between cohorts:** (C) Agathobacter rectalis, (D) Blautia_A wexlerae, (E) Fusicatenibacter saccharivorans, (F) Enterococcus_B faecium. Post hoc Dunn’s tests revealed significant enrichment patterns, depicted by lines above boxplots.

### The gut microbiome of ACLD patients represents an enriched reservoir of antimicrobial resistance

Antimicrobial resistance profiling was conducted on metagenomic assemblies to assess the prevalence of ARGs across study groups. Notably, ARG abundance was significantly higher in ACLF compared to other cohorts. These ARGs spanned 20 unique antimicrobial classes, including beta-lactams, aminoglycosides, glycopeptides, and tetracyclines (Figure 3A). HC and SC harboured a limited number of genes associated with tetracycline, sulphonamide, and macrolide-lincosamide-streptogramin within their resistome (Figure 3A). These may reflect intrinsic resistance genes that occur naturally across several taxa and contribute to baseline resistome levels, independent of antimicrobial exposure. However, a stepwise expansion of ARG burden was observed for multiple antimicrobial classes. For example, beta-lactam resistance genes were present in low abundance in HC and SC but were enriched in DC and ACLF. Similar trends were observed for quinolone, phenicol, and tetracycline resistance determinants, which were significantly enriched in DC and ACLF. This enrichment may be partially driven by clinical exposure to antimicrobials such as ciprofloxacin, norfloxacin, and doxycycline (Supplementary Table 1). Community-level differences were explored using NMDS based on Jaccard dissimilarities (Figure 3B). HC formed a tight cohesive cluster, whereas SC, DC, and ACLF displayed progressively greater dispersion with increasing disease severity. Testing for unequal dispersion using betadisper() confirmed heterogeneity (P = 0.003). To account for this, non-parametric ANOSIM confirmed significant differences between patient groups and HC (P=0.002).

**Figure 3.**
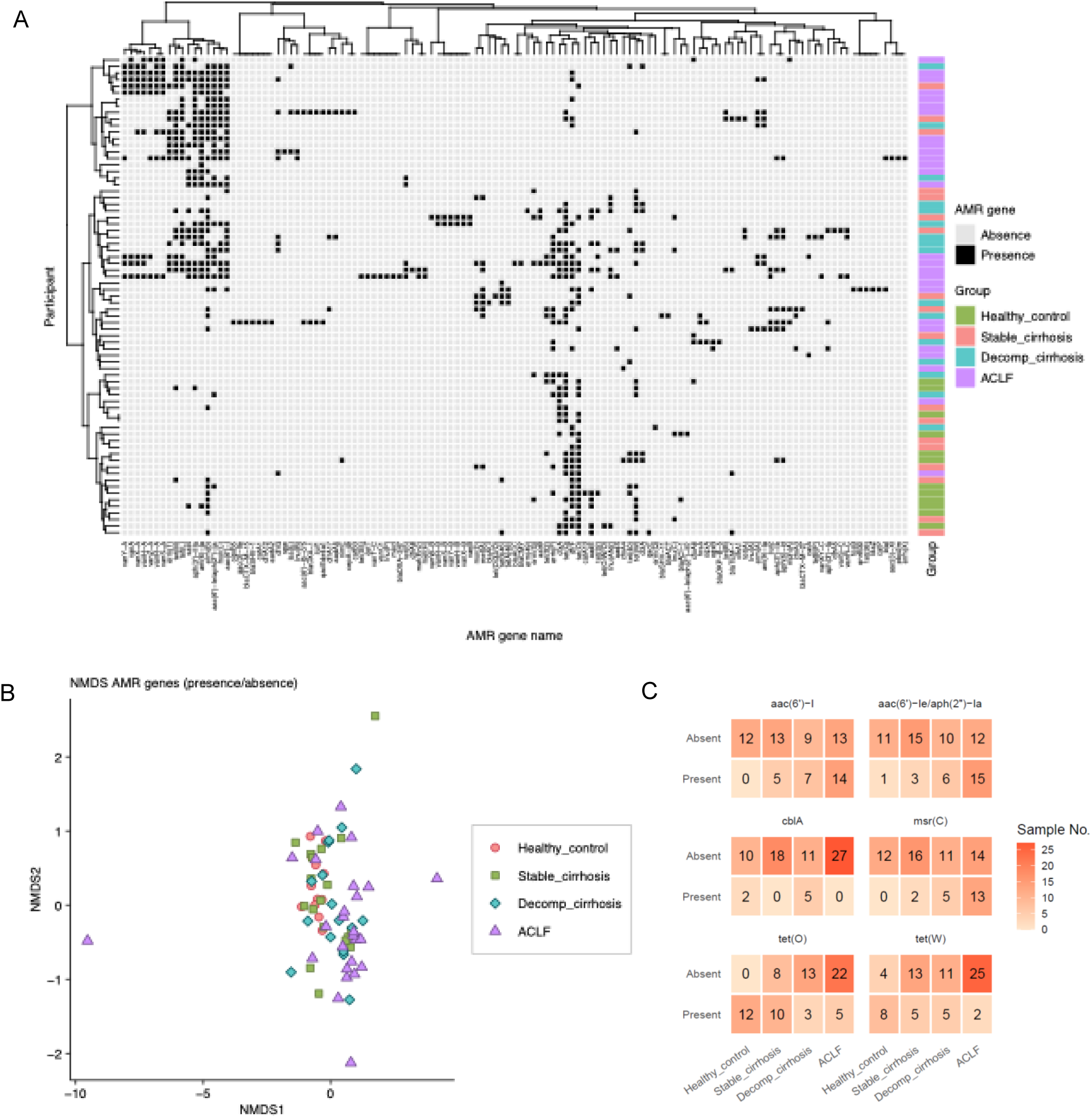
Distribution and clustering of antimicrobial resistance genes. **A)** Clustered heatmap showing presence (black) and absence (grey) of ARGs within cohort samples. **B)** NMDS plot based on Jaccard dissimilarities of ARG presence/ absence, displaying clustering patterns by disease severity. **C)** Differentially distributed ARGs across cohorts for*, aac(6’)-I, aac(6’)-Ie/aph(2’’)-Ia, cblA, msr(C), tet(O), and tet(W).* Numbers represent sample counts with ARGs present within each cohort.

At the gene level, differential distribution was assessed using Fisher’s exact test or Chi-squared test, depending on counts. Six ARGs showed statistically significant differences: *aac(6’)-Ie/aph(2’’)-Ia, tet(O), tet(W), msr(C), aac(6’)-I,* and *tet(L)* (Figure 3C). Of these genes, four genes (*tet(O), tet(W), msr(C)*, and *tet(L)*) were more frequently observed in HC. In contrast, aminoglycoside-modifying genes *aac(6’)-Ie/aph(2’’)-Ia* and *aac(6’)-I,* exhibited a stepwise increase in prevalence, enriched in DC and ACLF cohorts. Similar enrichment patterns were observed for ARGs within the glycopeptide class, including *vanA* and *vanB,* as well as their variants.

### Recovery of metagenome assembled genomes from clinical samples highlights strain-specific resistomes

Draft genomes were generated from the metagenomes by grouping contigs based on their shared properties, resulting in metagenome-assembled genomes (MAGs). In total, 389 MAGs were generated with completeness > 70 % and contamination < 10 %. Dereplication at 99.9% ANI yielded 382 nonredundant strain representative MAGs, excluding seven genomes identified as *E. faecium.* Completeness ranged from 70.07 % to 100 %, and contamination ranged from 0 % to 9.918 % (Figure 4A). The MAGs represented 10 phyla, of which Bacillota was the most abundant phylum (n=255), Actinomycota (n=54) and Bacteroidota (n=50). The most prevalent species within the dereplicated MAGs were *E. faecium* (n=22) followed by *B. longum* (n=17) (Figure 4B).

**Figure 4.**
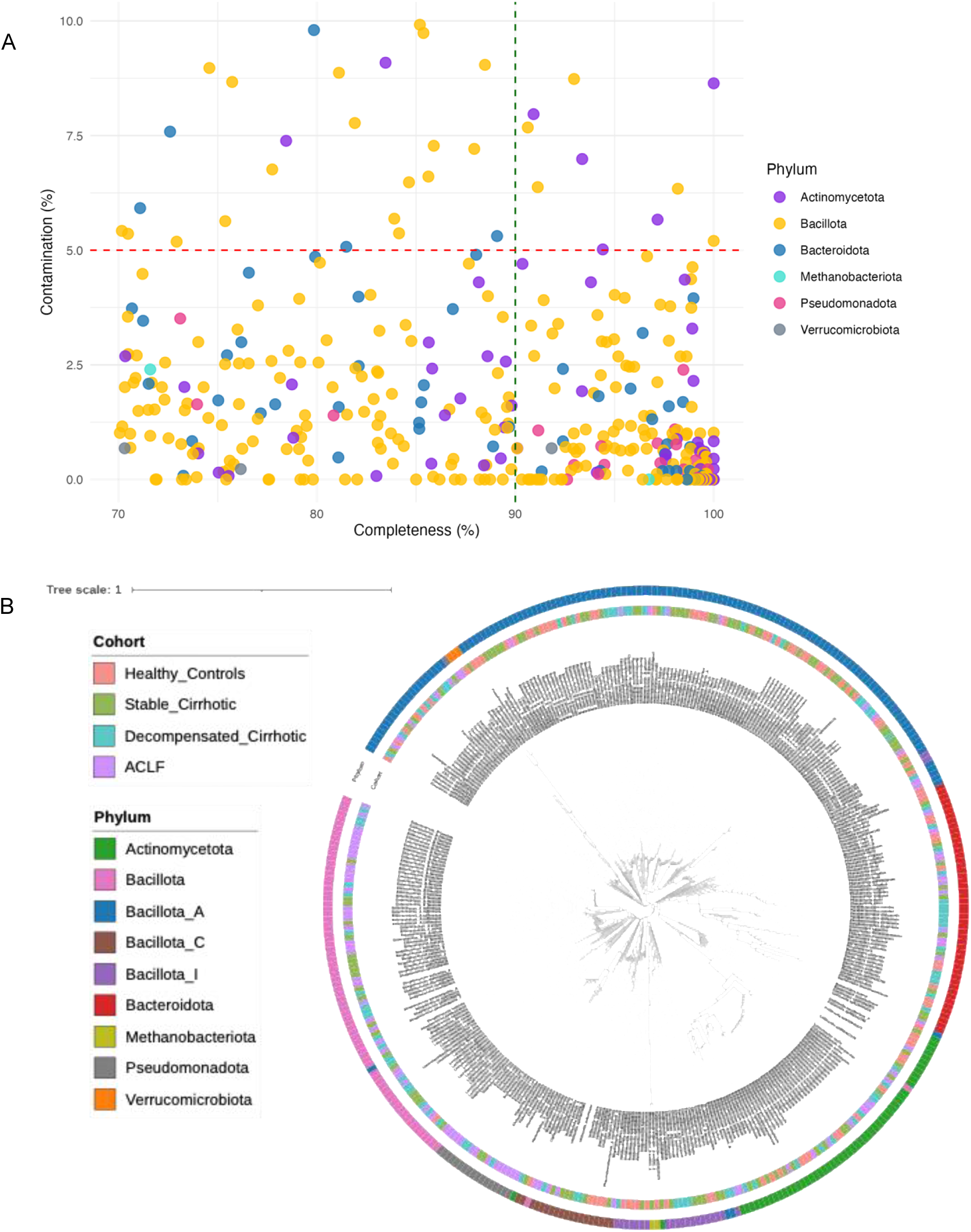
Phylogeny and quality assessment of metagenome-assembled genomes. **A)** Scatter plot of MAG quality showing completeness (x-axis) versus contamination (y-axis). High-quality MAG thresholds, as defined by MIMAG criteria (The Genome Standards Consortium et al., 2017) are indicated by dotted lines: green for completeness and red for contamination. Individual MAGs are color-coded by phylum.**B)** Strain level neighbour joining tree of bacterial strains (n=382) colour coded by cohort and phylum, with species IDs annotated in outer rings.

Given previous reports, including our own, linking *E. faecium* to advanced cirrhosis; we examined this species in greater detail. Twenty-eight MAGs were retrieved from SC, DC, and ACLF cohorts and placed within a midpoint-rooted maximum-likelihood tree, alongside reference genomes from Lebreton et al.(Lebreton *et al*., 2013) (Figure 5). Most *E. faecium* MAGs clustered tightly with A1 clade reference genomes, except for one MAG from a SC patient, which grouped within the A2 clade (Figure 5). MLST analysis revealed that A1 cluster MAGs belonged to clonal complex 17 (cc17), with predominant sequence types (STs) of ST78, ST80, ST117, and ST192, which are lineages previously associated with hospital-acquired transmission and multidrug resistance. Overall, these MAGs displayed extremely high similarity over 97 %, with several pairs sharing over 99.9 % ANI (strain-level cut-off) (Supplementary Table 2), suggesting potential strain sharing within patient groups, with persistence of *E. faecium* strains across individuals.

**Figure 5.**
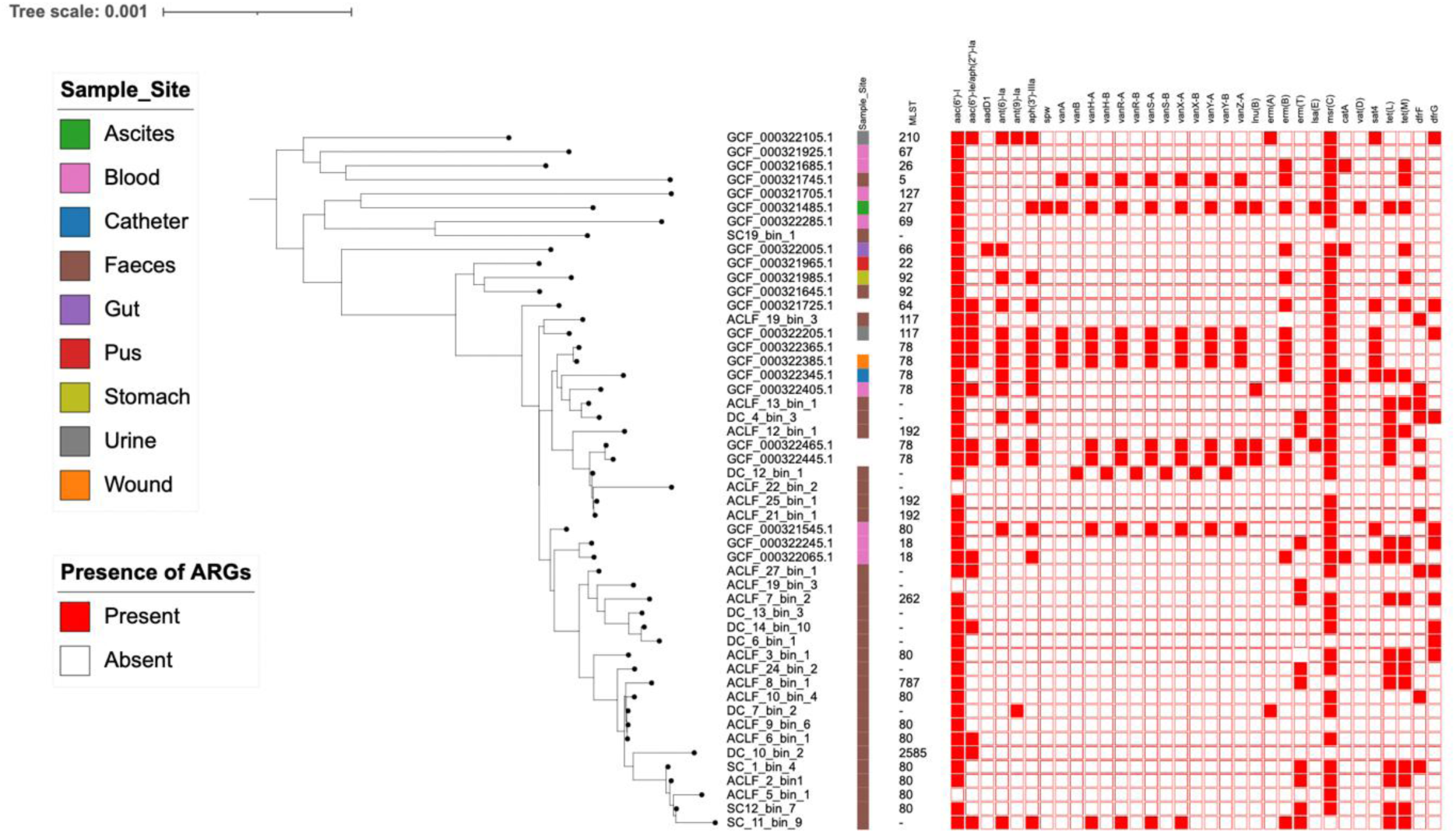
Phylogenetic tree of E. faecium MAGs and 22 reference genomes. A mid-rooted phylogenetic tree of 28 E. faecium MAGs and 22 reference genomes was constructed and visualised using iTOL v7. Sequence types from PubMLST and are indicated alongside each genome. Sample sites are denoted by coloured symbols. Presence/absence of ARGs is shown as a matrix alongside the tree (red = presence, white = absence).

Phylogenetic analysis of 28 *E. faecium* MAGs and 22 reference genomes revealed distinct patterns of ARG distribution. Vancomycin resistance genes (*van*) were largely absent from MAGs, including those within the A1 cluster associated with hospital-acquired infections, despite belonging to cc17 based on MLST (Figure 5). The macrolide/streptogramin resistance gene *msr(C)* was detected in nearly all genomes, absent in only eight MAGs. Aminoglycoside resistance gene *sat(6’)-I* was absent in three MAGs; however, *aac(6’)-Ie/aph(2”)-Ia, ant(6)-Ia,* and *apt(3”)-II* were consistently present in reference genomes but largely absent in MAGs. Notably, *van* and *aac(6’)-Ie/aph(2”)-Ia* were detected in the metagenomic dataset, coinciding with an increase in *E. faecium g*abundance, suggesting that resistance determinants may be associated with species expansion.

### Multiple ARGs are encoded on plasmids within the ACLD gut microbiome

Whilst MAGs provided species-specific insights into ARG presence, metagenomic binning does not capture plasmids, which are major contributors to ARG dissemination within bacterial hosts. To address this limitation and overcome the underrepresentation of plasmid-associated ARGs in the MAG analysis, plasmid contigs were identified within the metagenomes and profiled to examine the distribution of ARGs. Across the 76 faecal samples, 64 contained plasmid-assigned contigs that harboured ARGs (Figure 6a). The clustered heatmap of plasmid-associated ARGs revealed distinct resistance signatures across disease stages. HC plasmid contigs encoded a limited number of ARGs, such as β-lactam and tetracycline resistance determinants, without significant enrichment by disease stage. Several β-lactamase genes were detected across all groups, including *blaTEM-1*, *blaCTX-M-27*, *blaOXA-1*, and *blaSHV-11,* alongside tetracycline resistance genes *tet(O), tet(W), and tet(Q*), which are more frequently observed in HC and SC. In more advanced cirrhosis patient samples, increases in *erm* genes (conferring resistance to lincosamide/macrolide/streptogramin) and aminoglycoside resistance determinants were detected in DC and ALCF (Figure 6a).

**Figure 6.**
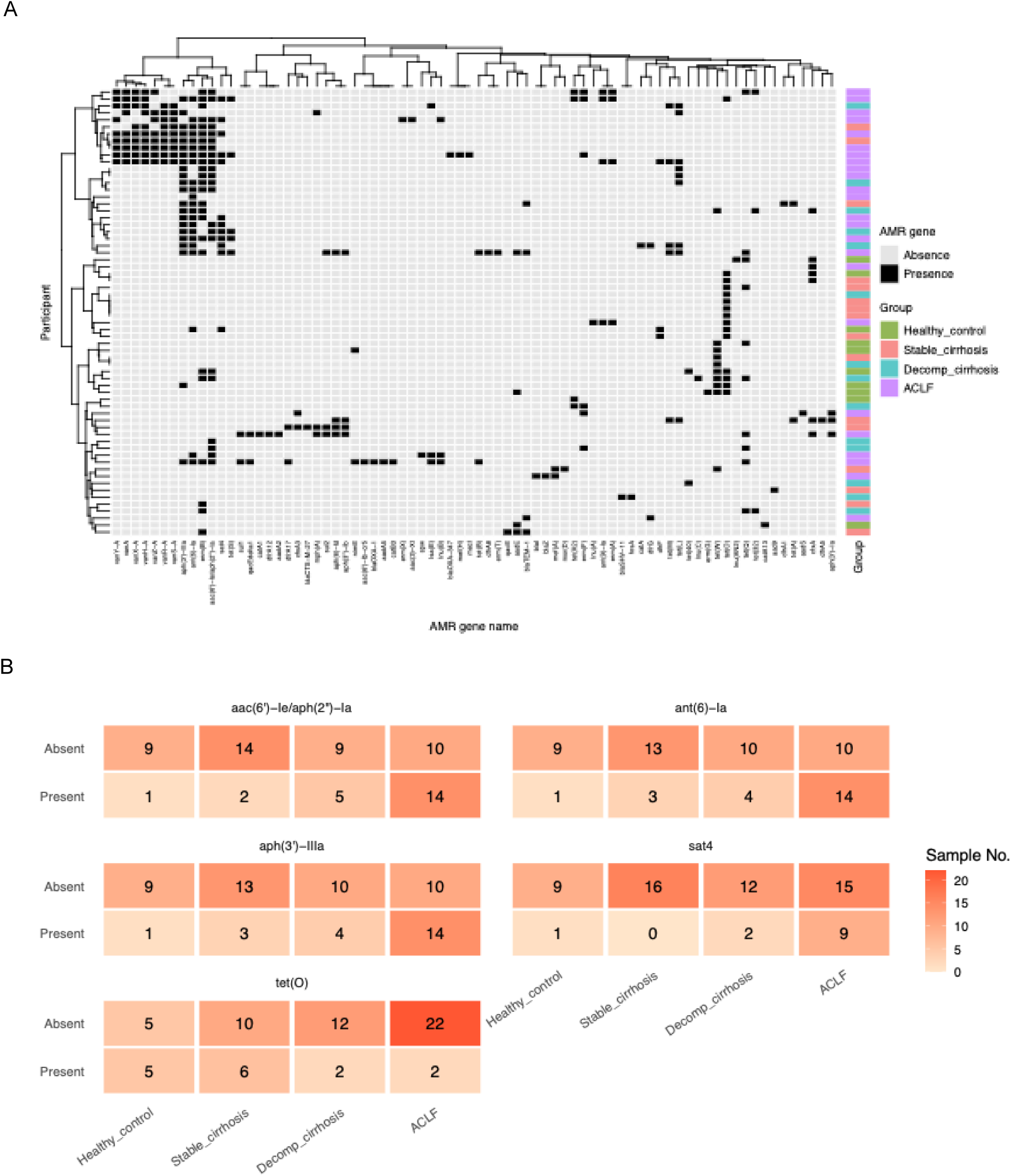
Clustered presence/absence heatmap of putative plasmid-associated ARGs grouped by drug class. **A)** Heatmap showing ARG distribution on plasmid-derived contigs identified by GeNomad across samples. ARGs grouped by antimicrobial class; black = presence, grey = absence. Group annotations denote participant categories: HC, SC, DC, ACLF. **B)** Statistically significant, differentially distributed ARGs across cohorts for aac(6’)-Ie/aph(2’’)-Ia”, ant(6)-Ia, aph(3’)-IIIa, tet(O), and sat4. Numbers represent sample counts with ARGs present on plasmid contigs.

Differential distribution analysis confirmed five ARGs with significant group-specific differences: *aac(6’)-Ie/aph(2’’)-Ia*, *ant(6)-Ia*, *aph(3’)-IIIa*, *sat4*, and *tet(O)* (Figure 6b). Aminoglycoside resistance genes, including *aac(6’)-Ie/aph(2’’)-Ia, ant(6)-Ia, aph(3’)-IIIa*, and *sat4*, were strongly enriched in ACLF patients, with the first three detected in 14 of 24 ACLF samples and *sat4* present in nine, highlighting a marked expansion of aminoglycoside resistance in advanced cirrhosis. In contrast, tetracycline resistance was more frequent in HC and SC (e.g., *tet(O)* in 5/15 HC and 6/18 SC) but largely absent in ACLF samples.

In metagenomes where *E. faecium* was present, there was an increased presence of ARGs, particularly glycopeptides and aminoglycoside resistance determinants (including streptothricin resistance), consistent with the intrinsic resistance profile of *E. faecium* (Figure 6a). Screening plasmid-derived contigs identified only one contig classified as an *E. faecium* plasmid carrying a *van* gene cluster, using ≥ 90% identity and coverage thresholds.

### Using MARTi (Metagenomic Analysis in Real Time) open-source software to profile metagenomes

We wanted to trial using a tool that could provide rapid real-time results in a clinical setting; therefore we supplemented the results gathered using traditional bioinformatic methodologies with *MARTi* (Metagenomic Analysis in Real Time). We applied MARTi to a subset of 35 samples to evaluate its potential for rapid, open-source metagenomics profiling, (Peel, *et al*., 2025). MARTi employs a read-based approach to classify sequencing data, enabling simultaneous taxonomic profiling, detection of ARGs, and attribution of ARGs to specific taxa. This read-level resolution is particularly advantageous for complex clinical samples, where accurate characterisation of both microbial diversity and resistance determinants is essential. Taxonomic profiling of the top 25 genera highlighted a progressive decline in diversity from SC, through to ACLF (Figure 7A). ACLF samples were dominated by opportunistic taxa, including *Escherichia* and *Enterococcus,* accompanied by marked depletion of keystone beneficial genera such as *Faecalibacterium, Blautia,* and *Anerostipes.* ARG analysis demonstrated broad resistance diversity across the subset, with aminoglycosides, glycopeptides and tetracyclines representing the top three drug classes (Figure 7B). These patterns likely reflect previous exposure to antibiotics (Supplementary Table 1). Four ACLF samples (ACLF6, ACLF7, ACLF13 and ALCF27) had very high ARG read counts (>5,000), highlighting a substantial resistance burden. Species-level attribution identified *E. faecium* and *E. coli* as the predominant ARG carriers (Figure 7C). Notably, *E. faecium* showed particularly high ARG loads in ACLF 6, ACLF7, ACLF 13, ACLD 21 and ACLF27, each exceeding 1,000 ARG reads.

**Figure 7.**
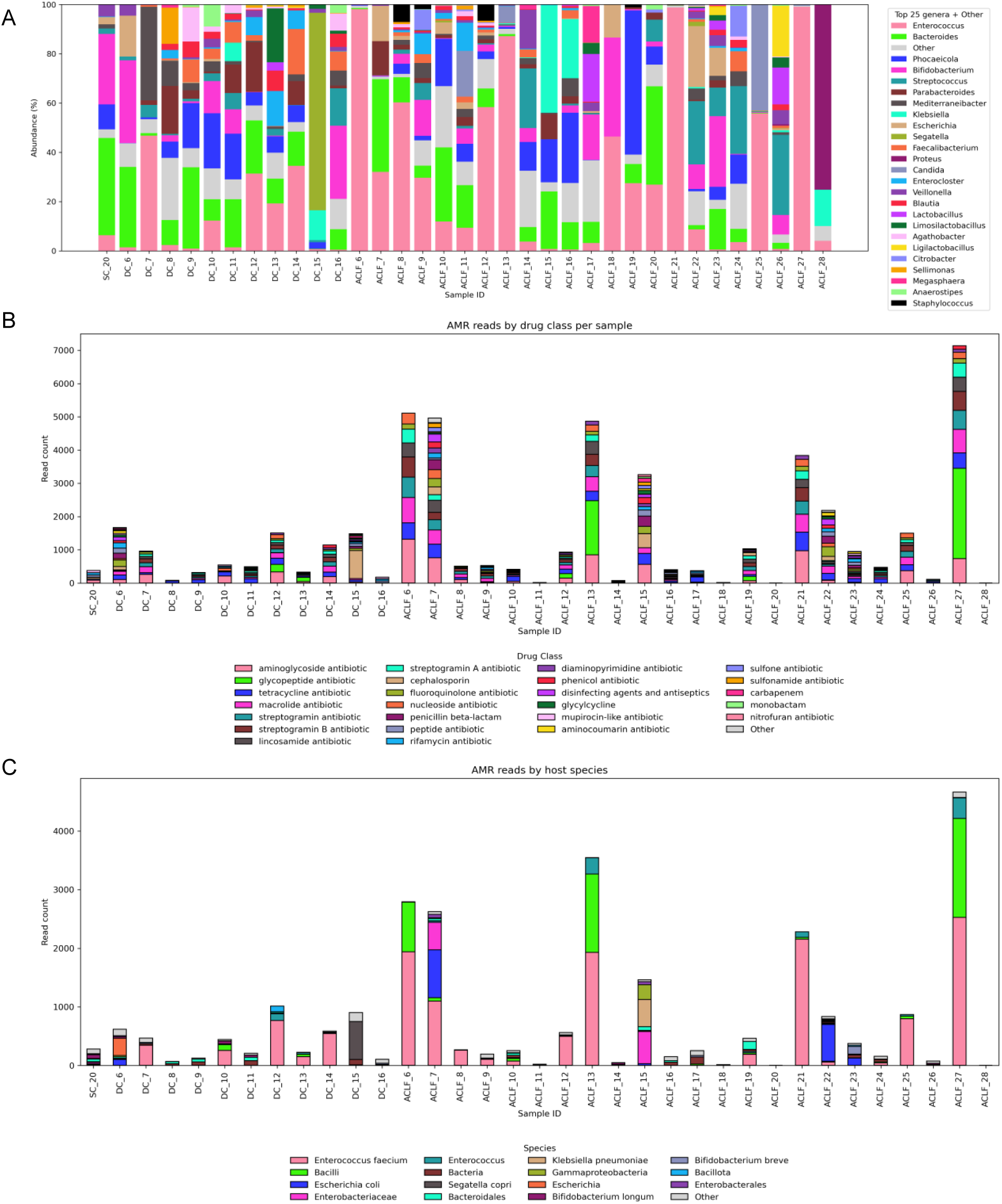
Comparative analysis of reads for AMR and taxonomic profiles produced by MARTi. **A)** Taxonomic relative abundance plot using LCA minimum abundance threshold of 0.1%. Top 25 taxa are shown, all others are collected under “Other”. **B)** AMR read count bar plot for top 25 drug classes. Read counts mapping to AMR genes were aggregated and grouped by their associated resistance drug class. Because a single AMR gene may have multiple associated mechanisms or classes, reads can be counted in more than one category**. C)** Potential host taxa of AMR genes for top 15 bacterial taxa. Host taxa of AMR genes identified using walkout analysis within MARTi using a minimum threshold of 10%, reporting a potential host of an AMR gene if at least 10% of the AMR-matching reads were assigned to that taxon.

## Discussion

This study used ONT long-read metagenomic sequencing to characterise the gut microbiome and resistome in patients with ACLD, providing insights into microbial community structure and AMR spanning different ACLD severities. The untargeted nature of metagenomic sequencing enabled strain-level identification without prior assumptions, overcoming the limitations of culture-based diagnostics (Govender *et al*., 2021). Historically, such approaches required advanced bioinformatic expertise; however, the emergence of plug-and-play bioinformatic workflows has significantly improved the accessibility and usability of metagenomic sequencing, with data analysis now integrated into intuitive online platforms that generate clear and concise outputs (Fan, Huang and Chorlton, 2021; Kelliher *et al*., 2024).

Metagenomic profiling of the ACLD patient faecal samples revealed striking and progressive alterations in microbial community structure across disease severities. Diversity indices (Chao1, Shannon, and Inverse Simpson) revealed a stepwise decline from HC to ACLF, supported by PERMANOVA and NMDS clustering. This progressive loss of richness and evenness coincided with a compositional shift from beneficial taxa, such as butyrate-producing taxa like *Faecalibacterium*, *Blautia*, and *Agathobacter*, towards opportunistic pathogens, most notably *Enterococcus* and *Escherichia*. These changes were most pronounced in ACLF, where *E*. *faecium* frequently dominated (>50% relative abundance in multiple samples), especially among patients with more advanced cirrhosis and a history of multiple antibiotic treatments. This expansion reflects disruption of the gastrointestinal ecosystem, consistent with previous reports. It is likely linked to reduced colonisation resistance and compromised mucosal integrity, factors that facilitate pathogen overgrowth and translocation (Solé *et al*., 2021; Dalby *et al*., 2025; Tyc *et al*., 2025). In contrast, *E. faecium* was not detected in HC samples. While this cross-sectional dataset cannot determine the source of colonisation, the absence of *E. faecium* in healthy participants who served as negative controls in this study suggests that healthcare staff are unlikely to act as reservoirs within this cohort, or indeed are themselves colonised with nosocomial strains. However, hospital-associated acquisition in and between patients remains possible from other hospital reservoirs, such as hospital sinks, medical equipment, bedding, and the air (Kanamori, Rutala and Weber, 2017; Dalton *et al*., 2020, 2020).

The depletion of beneficial, particularly butyrate-producing, taxa is both ecologically and clinically significant. Microbes, such as *Faecalibacterium, Roseburia, Ruminococcus,* and *Agathobacter*, play key roles in gut homeostasis by fermenting nondigestible carbohydrates into short-chain fatty acids, such as butyrate (Hodgkinson *et al*., 2023). Butyrate supports epithelial barrier function, modulates immune responses, and exerts anti-inflammatory effects, and its loss may therefore exacerbate (Plöger *et al*., 2012) gut permeability and inflammation, creating conditions that favour opportunistic pathogens such as *Enterococcus*, and facilitating bacterial translocation into the bloodstream (Acharya and Bajaj, 2019). Loss of these beneficial taxa is a common feature of ACLD, often linked to altered bile acid metabolism, repeated exposure to broad spectrum antibiotics, and dietary changes (Dalby *et al*., 2025). A caveat to this observation was of *Bifidobacterium* in DC and ACLF cohorts. This genus is usually susceptible to broad spectrum antibiotics, displaying high antimicrobial sensitivity (Zimmermann and Curtis, 2019); however, *Bifidobacterium* persisted, albeit at lower abundances than in HC, in advanced disease stages, potentially reflecting the influence of lactulose therapy, which has prebiotic effects and promotes its growth, which is an observation supported by cohort metadata and previous reports.

The overgrowth of *E. faecium* was significant in both DC and ACLF patients, consistent with previous findings (Dalby *et al*., 2025; Tyc *et al*., 2025). Genomic analysis revealed high strain diversity (97 – 99.9 % ANI) with 27 of 28 genomes clustering within the A1 clade, a lineage commonly associated with hospital-acquired infections. This clade links to nosocomial persistence, biofilm formation, and accumulation of multiple ARGs (Wei, Palacios Araya and Palmer, 2024), reflecting strong selective pressures within the cirrhotic gut, driven by frequent antibiotic exposure, compromised mucosal epithelial barriers, and immune dysfunction, which collectively favour the expansion of multidrug-resistant, *E. faecium* lineages, and possible strain sharing between patients. In a prospective cohort of ACLD patients, circulating *E. faecium* DNA was associated with heightened systemic inflammation and renal dysfunction, proposing *E. faecium* translocation as a key consequence of gut barrier failure in ACLF and providing further clinical context for microbiome signatures centred on this pathobiont (Tyc *et al*., 2025). However, it is also possible that the strains represent endogenous populations of *E. faecium,* which exploit the selective conditions presented as they evolve and expand within the host (Hourigan *et al*., 2024). The genomic diversity observed across MAGs indicates potential for independent within-host emergence of these particular strains under similar selective pressures, rather than from a single nosocomial source.

In keeping with the multidrug-resistant characteristic of A1 and cc17 lineages, vancomycin resistance genes (*vanA, vanB*) were enriched in the metagenomes from DC and ACLF patients. The *van* gene complexes, typically carried on MGEs, mediate high-level vancomycin resistance and facilitate HGT across taxa, and have been attributed to the overprescription and reliance on vancomycin to treat infections (Almeida-Santos *et al*., 2025). Their presence aligns with global concerns over the emergence of vancomycin-resistant *E. faecium,* recognised by the WHO as a critical priority pathogen (*WHO Bacterial Priority Pathogens List 2024: Bacterial Pathogens of Public Health Importance, to Guide Research, Development, and Strategies to Prevent and Control Antimicrobial Resistance*. 1st ed, 2024). Further genomic analysis confirmed the predominance of cc17-associated A1 clade MAGs; however, *van* genes were largely plasmid-borne and absent from MAG assemblies, reflecting both their extrachromosomal localisation and shallow sequencing depth in this study. The ONT R9 chemistry, though valuable for long-read sequencing, has lower accuracy than newer versions. Combined with limited metagenomic depth, this likely reduced strain-level resolution and plasmid recovery, contributing to the apparent underrepresentation of some ARGs in MAG-based analyses.

Notably, vancomycin-resistant *E. faecium* is also multidrug resistant and is often co-resistant to gentamicin (Radford-Smith and Anthony, 2025). As highlighted above, aminoglycoside resistance genes were detected within the wider resistome; however, the bifunctional aminoglycoside acetyltransferase gene *AAC(6’)-Ie-APH(2’’)-Ia* could not be directly linked to *E. faecium*. This gene was frequently identified on plasmid-like contigs and significantly enriched in ACLF faeces. The current dataset lacked sufficient resolution to confidently associate plasmids with their bacterial hosts, constraining the ability to resolve MGE dynamics within the microbiome. Whilst specific bacterial hosts could not be conclusively linked to these ARGs, the increased prevalence correlates with the abundance of *E. faecium*, supporting its role as a likely reservoir of glycopeptide resistance. The co-localisation of multiple ARGs on MGEs is particularly concerning, as it underscores the potential for co-transfer of multidrug resistance, increasing dissemination within the gut microbiome, and limiting effective treatment options for severely ill patients, such as those with ACLF.

Taking a broader look at the total microbiome resistome, AMR analysis revealed a clear correlation between ACLD severity and the escalation of resistance burden and has been recently reported to affect not only the gut microbiome but also the oral cavity (Lee *et al*., 2025). ARGs spanned multiple classes, including β-lactams, aminoglycosides, glycopeptides, and tetracyclines. While HC and SC harboured limited ARGs, ACLF samples exhibited extensive resistomes, with aminoglycoside-modifying genes (*aac(6’)-Ie/aph(2’’)-Ia*) strongly enriched. Plasmid-associated ARGs were common in advanced cirrhosis samples, highlighting the potential for HGT and rapid dissemination of multidrug resistance within the gut ecosystem. These resistome patterns, combined with impaired colonisation resistance due to depletion of beneficial taxa, align with clinical metadata showing frequent broad-spectrum antibiotic exposure in DC and ACLF patients. In DC and ACLF patients, broad-spectrum antibiotics such as piperacillin/tazobactam, meropenem and rifaximin were frequently administered. These antibiotics are often prescribed, prior to infection confirmation, because culture-based diagnostics can take up to a week to isolate and identify causative pathogens (Fernández *et al*., 2021). Such selective pressures likely facilitated the expansion of resistant opportunistic pathogens, including *E. coli*, *K. pneumoniae* and *E. faecium*, while depleting commensal populations that normally confer colonisation resistance.

In addition to traditional methodologies used to survey the microbiomes and resistomes of patients with ACLD, we utilised MARTi, a real-time, open-source metagenomic analysis platform (Peel *et al*., 2025), which provides a comprehensive read-based taxonomic profiling and host attributions of ARGs. The platform’s user-friendly interface greatly enhances accessibility, which is crucial for wider adoption in clinical settings, where bioinformatic training is often limited by complexity and expense (Trivett, Darby and Oyebode, 2025). It’s ability to analyse reads progressively, updating results as sequencing continues, further enhances its potential as a diagnostic tool. While MARTi relies on reference databases and may therefore miss rare or complex taxa and ARGs, and has limited capacity for functional profiling beyond ARG detection, these gaps can be addressed by supplementing the analysis with conventional metagenomic approaches. Nevertheless, MARTi’s accessible interface makes it well-suited for clinical applications, providing rapid, actionable insights where computationally intensive traditional pipelines may be impractical. These findings demonstrate the translational potential of metagenomic sequencing coupled with easy-to-use analysis platforms in clinical microbiology, enabling rapid pathogen detection and AMR surveillance, particularly for severely ill patient groups, such as those with ACLF.

While these findings highlight distinct microbial and resistome alterations in ACLD, the cross-sectional study design precluded the assessment of temporal dynamics, preventing evaluation of how microbial and resistome profiles evolve during disease progression and treatment. Future work should address these limitations by employing higher accuracy sequencing chemistries, deeper metagenomic coverage, and longitudinal sampling. These refinements would enhance MGE and plasmid recovery, enable robust plasmid-host interactions, and provide clear insights into the evolution of ARGs over time as ACLD progresses. Incorporating these capabilities into workflows like MARTi would also further enhance their translational and clinical applicability and diagnostic value. Together, such improvements would help define strain-specific microbial and resistome signatures of ACLD, informing therapeutic interventions and infection control, including surveillance strategies in this high-risk population.

To conclude, long-read metagenomic sequencing demonstrates substantial potential for characterising both the microbiome and resistome in patients with ACLD who are at very high risk of infections that require hospitalisation. The culture-independent approach provides a comprehensive view of the gut ecosystem, simultaneously identifying pathogenic taxa, community composition, and circulating ARGs. The ability to detect clinically relevant species such as *E. faecium* and their associated ARGs directly from metagenomic data highlights its value as a diagnostic and surveillance tool. The findings also highlight how chronic antibiotic exposure and microbiome disruption in ACLD create selective pressures that promote the expansion of multidrug-resistant pathogens such as *E. faecium*. Integrating metagenomic approaches, particularly through user-friendly, real-time analysis platforms like MARTi, into clinical workflows could support rapid pathogen detection, guide antimicrobial stewardship, and improve therapeutic decision-making, ultimately enhancing outcomes for ACLD patients at high risk of infection.

## Acknowledgements

We are grateful to all the patient participants and healthy volunteers for agreeing to take part in this study, and to the clinical and liver research teams at King’s College Hospital for facilitating recruitment, collecting metadata and sample collection.

## Author contributions

Conceptualisation: LJH, VCP, RL; Funding acquisition & supervision: LJH; Methodology & sequencing: DH; Sample processing: RK, AAG. Data analysis: MJD, NP, HT; QC, data processing, metagenomic assemblies, and data visualisation: HT; Clinical metadata curation: MM & GH; Clinical oversight & cohort design: VCP. Writing - original draft: HT, LJH; Writing - review & editing: All authors contributed to reviewing and editing the manuscript. All authors approved the final version.

## Ethical statement

The study was granted ethics approval by the national research ethics committee (12/LO/1417) and the local research and development department (KCH12-126) and performed in accordance with the Declaration of Helsinki. Patient participants, or their family nominee as consultees in the case of lack of capacity, provided written informed consent within 48 hours of presentation. Patients were managed according to standard evidence-based protocols and guidelines.

## Data summary

The metagenomic sequencing raw reads are deposited in NCBI SRA under project PRJNA1377628.

